## Supplementary data for "Pluripotency factors determine gene expression repertoire at zygotic genome activation"

### Supplementary Figures and legends

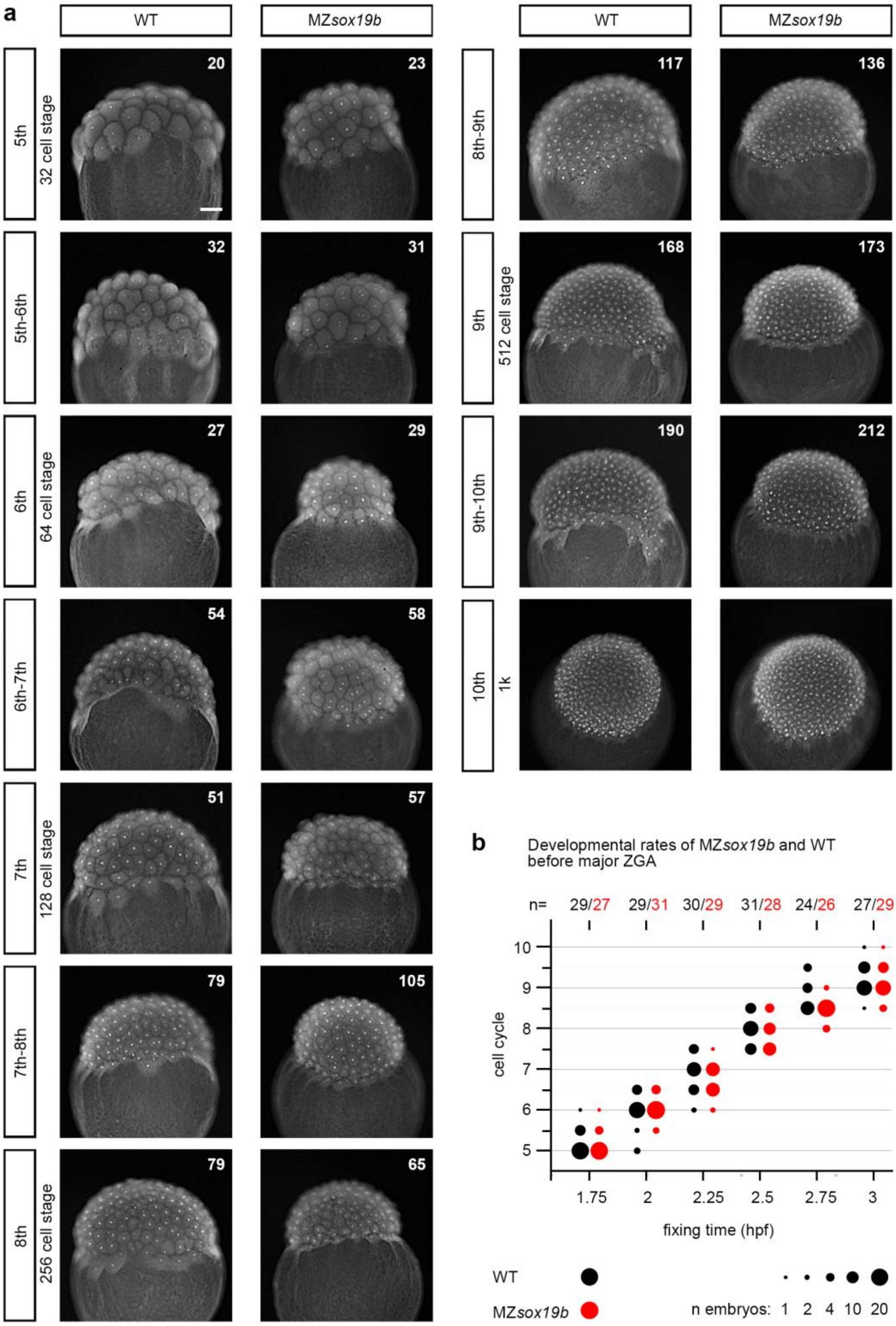

**Figure S1: Supporting information for the main Figure 1. MZ*sox19b* embryos are not delayed before MBT**. Wild-type (WT) and MZ*sox19b* eggs were *in-vitro* fertilized at the same time and kept at 28.5°C. 30~35 embryos from each genotype were fixed every 15 minutes starting from 1.75 hpf (5th cell cycle, 32 cell stage) till 3hpf (10th cell cycle, 1K stage). Fixed embryos were permeabilized and stained with 0.025µM SYTOX green nuclear stain. The images of all embryos were taken and used to estimate the cell cycle phase. **a.** Representative images of the embryos at the indicated cell cycles in two genotypes. The embryos with metaphase to anaphase nuclei were scored as intermediate between the cycles (i.e. cycle 5-6, 6-7 etc.) Numbers of scored nuclei on the top right corner. **b.** Pre-MBT synchronous cell cycles have the same length in the WT and MZ*sox19b* embryos. The quantification of representative experiment out of two. Numbers of embryos scored for each genotype at each time point are indicated above the graph. Scale bar: 100 µm.

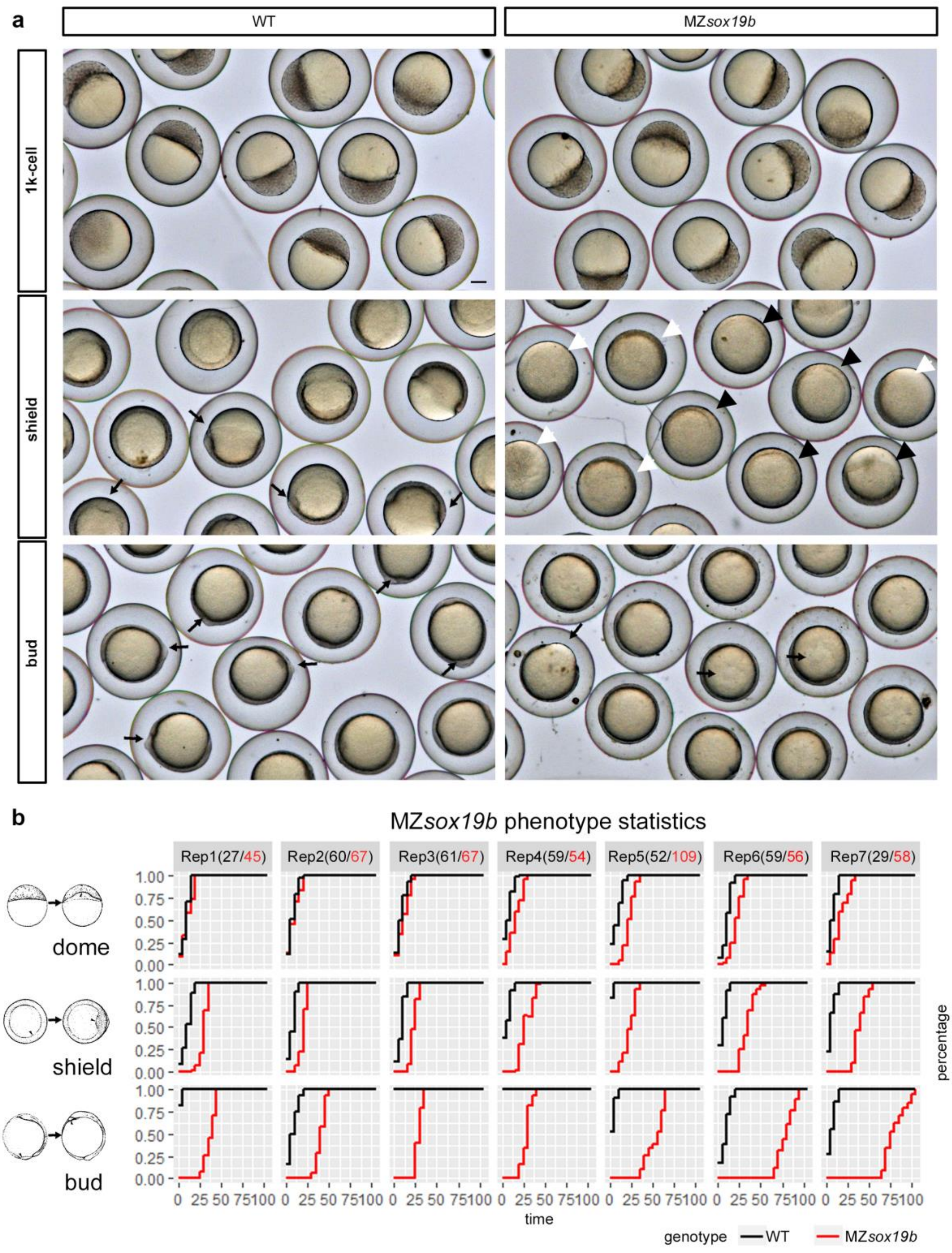

**Figure S2: Supporting information for the main Figure 1. *MZsox19b* embryos are delayed in gastrulation. a-b.** The embryos which were obtained from natural crosses in mass crossing cages. The fish were allowed to lay eggs for 15 minutes, then embryos were collected and let to develop at 28.5°C. **a**. Group pictures of the WT and MZ*sox19b* embryos at the indicated stages. 1k: no difference. Shield: Involution of mesendodermal layer marks the gastrulation onset, which occurs between 5.3 and 5.7 hours post fertilization (hpf) in the WT, and is followed by shield formation at 6 hpf at the dorsal side. Shield structure is visible in the WT embryos (black arrows in the WT). In the population of MZ*sox19b* embryos, mesendoderm only starts to involute: it is visible as an equatorial ring in some of the embryos (black arrowheads), but not at the others (white arrowheads). Bud: gastrulation ended in the WT, the yolk is completely covered by cell layer, tail bud is formed at the posterior side (black arrows in the WT show tail buds). MZ*sox19b* embryos are still undergoing gastrulation (black arrows show the free yolk). Scale bar: 200 µm. **b**. The developmental rates of the WT and MZ*sox19b* embryos was measured by scoring the time of appearance of developmental landmarks of three stages: dome (4.3 hpf), shield (6 hpf) and bud (10 hpf), as shown at the left. The stages were scored every 5 min. The results of 7 independent crosses from two generations of fish are shown; time in minutes, n embryos above the graph. X-axis: time in minutes; Y-axis: percent of embryos which reached the indicated stage (1 – 100%). Stages by Kimmel. Numbers of embryos per experiment on top of the graph.

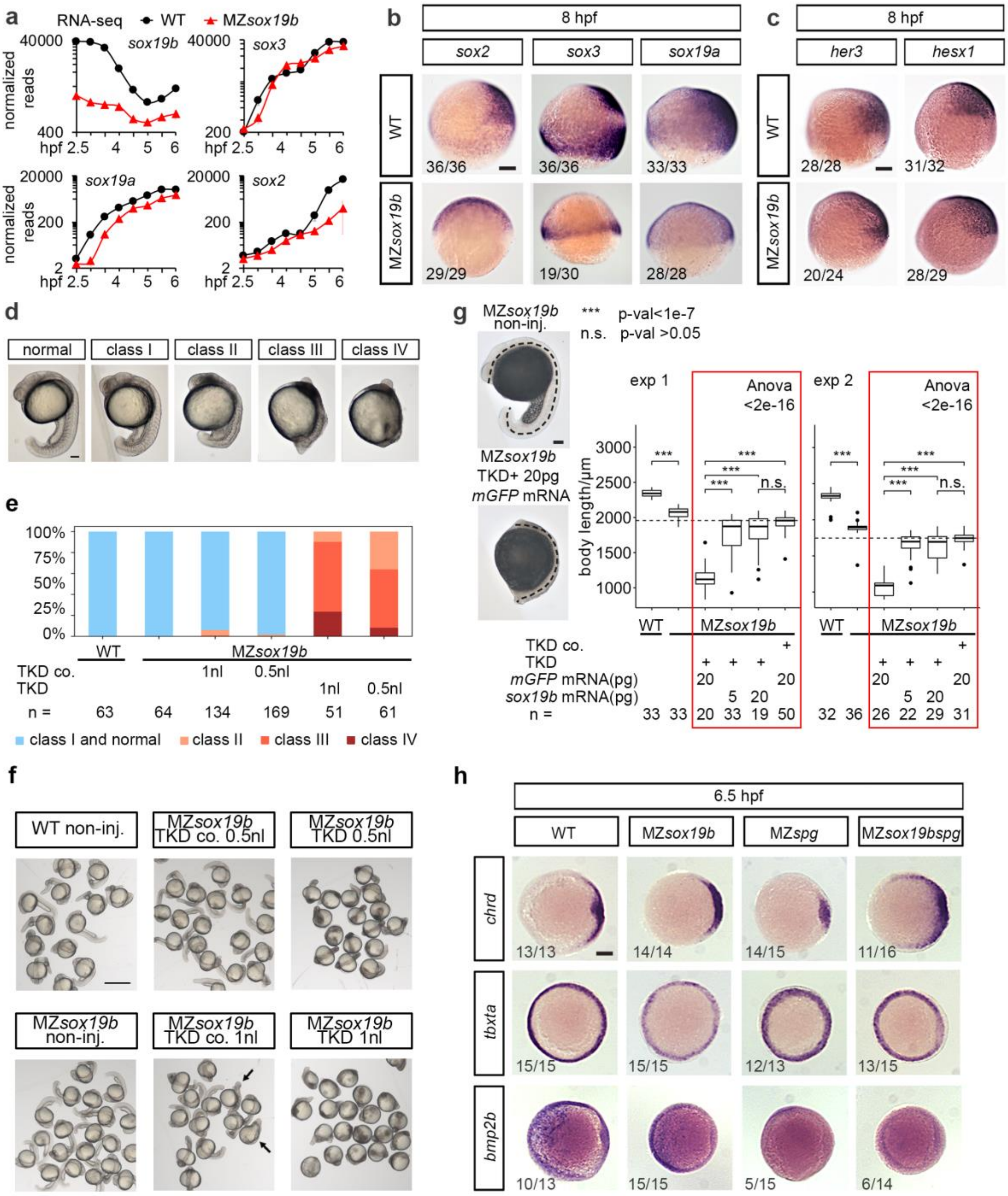

**Figure S3: Supporting information for the main Figure 1(a-g) and the main Figure 2 (g). a-c.** Zygotic SoxB1 family members Sox19a, Sox3 and Sox2 are present and functional in MZsox19b mutant. **a.** mRNA levels of *sox19b*, *sox19a*, *sox2* and *sox3* during development from 2.5 hpf to 6 hpf in the WT (black) and MZ*sox19b* (red), shown in the logarithmic scale. Note, that the levels of *sox19b* non-functional RNA in MZ*sox19b* are strongly reduced already at 2.5 hpf, due to non-sense RNA-mediated decay. Zygotic expression of *sox3*, *sox19a* and *sox2* in MZ*sox19b* starts with a delay after ZGA (3 hpf). Expression of the earliest zygotic SoxB1 members, *sox3* and *sox19a* expression recovers close to the WT levels with different dynamics. Expression of *sox2* is still strongly reduced at 6 hpf. RNA-seq, samples were collected in 30 min intervals (see Fig. S4a). **b.** Developmental delay was reflected in

the molecular patterning of the MZ*sox19b* mutants. *In situ* hybridization for *sox2, sox3 and sox19a*, 8 hpf, lateral views, dorsal to the right.
Note, that, at 8 hpf in the WT, *sox2*, *sox3* and *sox19a* became restricted to the prospective neuroectoderm; *sox3* is also expressed in the equatorial region of the embryo (mesodermal ring). In MZ*sox19b*, *sox2* and *sox19a* retained earlier ubiquitous pattern (compare to Fig. 1d in the main text). *sox3* was restricted to mesoderm, in the majority of MZ*sox19b* embryos (19/30 embryos, as shown in the figure), in 11 embryos *sox3* staining was ubiquitous (as in Fig. 1d, main text). **c**. Direct SoxB1 target genes *her3* and *hesx1*[1], are expressed in MZ*sox19b* mutants at 8 hpf. In situ hybridization, lateral views, dorsal to the right. Scale bar: 100 µm in (b-c). **d-f**. Morpholino titration experiment. To generate the triple knockdown of zygotic SoxB1 genes in MZ*sox19b* mutants, 1-cell stage MZ*sox19b* embryos were microinjected with Sox2, Sox3 and Sox19a morpholino mix (TKD mix, see Methods). In TKDco mix, Sox19a morpholinos were replaced with Sox19b morpholinos, so that only three out of four SoxB1 genes were blocked upon injection. TKDco was used to control for non-specific effects of TKD. To investigate the dose-dependent effects of the morpholinos, we injected 1 nl (5.4 ng) of TKD or TKDco per MZ*sox19b* embryo, or 50% of this (0.5 nl, 2.7ng). **d.** Phenotypic classes at 19 hpf, MZ*sox19b*. Injection of TKD in both amounts resulted in the range of phenotypes II-IV in two experiments. The “middle severity” phenotype (class III) was similar to previously published QKD phenotype in the WT, which was considered to be a complete SoxB1 knockdown[1]. The TKDco injection into MZ*sox19b* resulted in class I embryos, which were somewhat shorter than non-injected (shown as “normal”), indicating non-specific effects of Morpholino injections. Scale bar: 100 µm. **e**. Pooled statistics of two independent experiments. **f**. Group phenotypes at 19 hpf. Arrows show the embryos with strongest non-specific axial defects in 1 nl TKDco injected group, which were scored as “class II” (less than 5% in 1 nl injections) in (d). Scale bar: 500 µm. **g**. Triple Sox19b, Sox2 and Sox3 knockdown in MZ*sox19b* embryos (TKD) can be completely rescued by Sox19b mRNA. 1-cell stage MZ*sox19b* embryos were injected with TKD + control mRNA, TKD + 5 pg *sox19b* mRNA, TKD + 20 pg *sox19b* mRNA, or TKDco + control mRNA, as indicated. Non-injected WT and MZ*sox19b* embryos were collected at the same time and served as additional stage controls. We scored the live phenotypes at 19 hpf (Fig. 2c in the main text), then fixed all the embryos, took the lateral images and draw the midline from head to tail, as shown in the figure. The lengths of the body axes were measured in ImageJ. 1- way Anova and Tukey-Kramer test were performed to compare four injected samples (red frames at the right part of the graphs). The body length of TKD + control mRNA injected embryos was at least 1.5 times shorter than of the TKD + Sox19b mRNA-injected embryos, the difference was highly significant (1-way Anova $<2e-16$, p-val$<1e-7$ for all shown pairwise differences in Tukey-Kramer test). The differences between TKDco + control mRNA and TKD + *sox19b* mRNA-injected embryos were non- significant (p-val $>0.05$ in Tukey-Kramer test), and we therefore concluded that the rescue was complete. As expected, in both experiments the body length of the WT was longer than of MZ*sox19b* (left part of the graph, p-val = 3.7412e-26 in exp.1 and p-val = 2.2150e-25 in exp.2 in 2- tailed Student t-test). We also estimated non-specific morpholino effects, by comparing the body lengths of TKDco + control - injected MZ*sox19b* embryos with non-injected MZ*sox19b*. The difference was significant (p-val = 2.6503e-05 in exp.1 and p-val = 4.0969e-07 in exp.2 in 2- tailed Student t-test). n-number of embryos. Dotted line shows the mean value for TKDco + control mRNA – injected embryos. Scale bar: 100 µm. n=2 biologically independent experiments. The lower border, middle line and upper border of the box plot correspond to 0.25, 0.5

(median) and 0.75 quantile, respectively. **h.** MZ*sox19bspg* double mutants are more dorsalized than MZ*spg* and MZ*sox19b* mutants. *In-situ* hybridization with the probes for dorsal (*chrd*), pan-mesodermal (*tbxta*) and ventral (*bmp2b*) transcripts, animal view, dorsal to the right. Note that *chrd* is radially expanded in MZ*sox19bspg* compared to all other genotypes, while *bmp2b* staining is reduced in both MZ*sox19bspg* and MZ*spg*. Scale bar: 100 µm. Source data for the panel a are provided as a Source Data file.

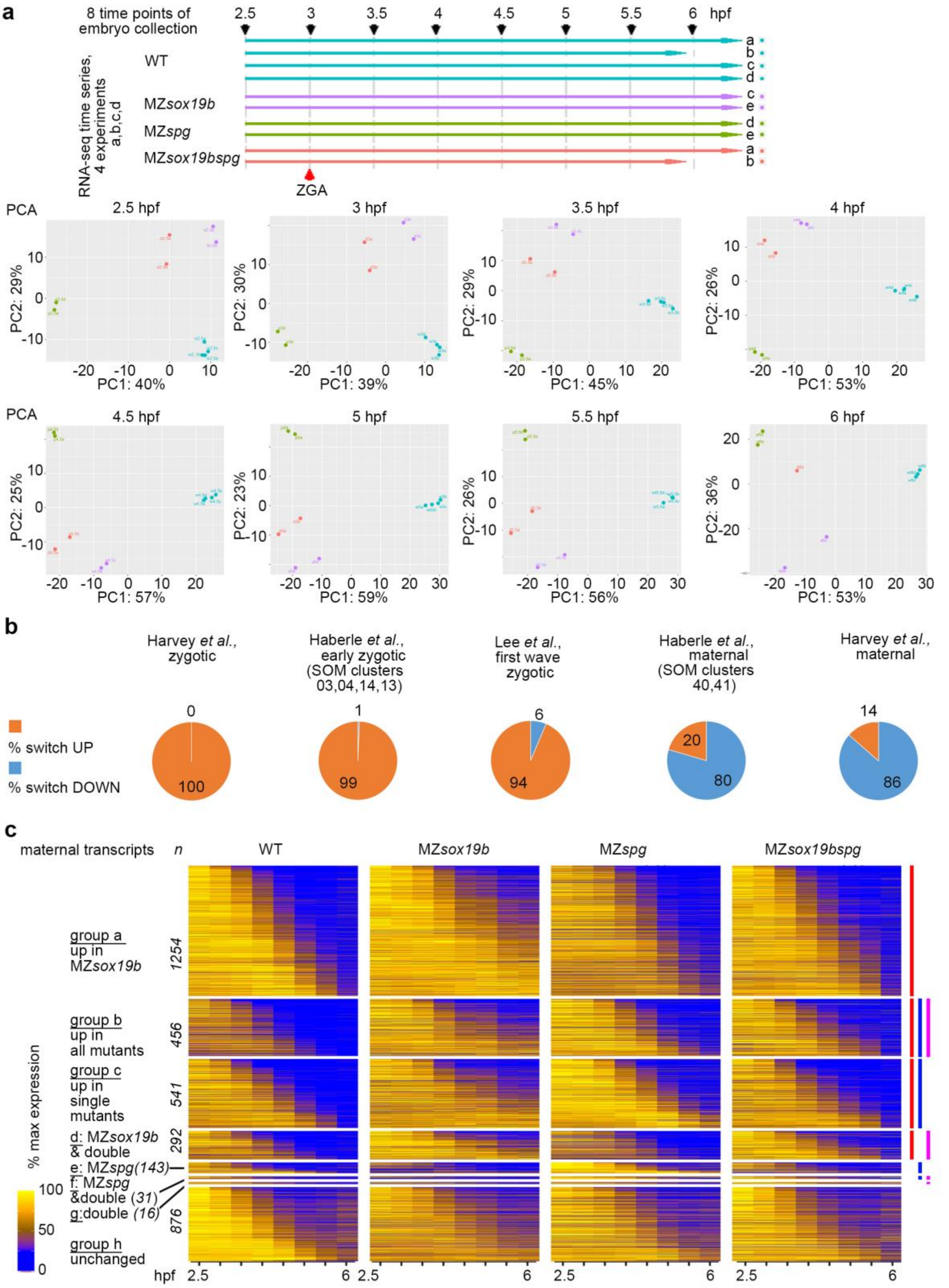

**Figure S4: Zygotic transcription and maternal RNA degradation are non-additively delayed in single and double mutants (supporting information for the main Fig. 3) a.** RNA-seq time curves, experimental setup and agreement between the replicates. Top: Five experiments, a-e, were performed at five different days. Embryos from two genotypes were collected in parallel in each of the experiments. In the experiments a, c, d, e the embryos were collected at 2.5, 3, 3.5, 4, 4.5, 5, 5.5 and

6 hpf. In the experiment b the embryos were collected at 2.5, 3, 3.5, 4, 4.5, 5, and 5.5 hpf. Bottom: Principle component analysis for each time point (Deseq2). **b.** Switching "UP" and "DOWN" transcripts match zygotic genes and maternal genes from previous studies: Harvey et al, 2013[2], Haberle et al., 2014[3], Lee et al., 2013[4]**.** The percentages of WT "switch UP" and "switch DOWN" genes are shown. The genes are listed in Dataset S1. **c**. Maternal RNA degradation is non-additively delayed in single and double mutants. Heatmap of all 3609 maternal transcripts in the indicated genotypes, grouped by regulation in one or several mutants. Color lines - upregulation in one of the mutant genotypes (red line - MZ*sox19b*, blue line – MZ*spg*, purple line - MZ*sox19bspg*) at the right. n- number of transcripts per group. 2733 transcripts were upregulated in one or several mutants, 876 transcripts were not (group h, unchanged). Dataset S1 lists all the transcripts and timepoints. Source data for the panels a,c are provided as a Source Data file. Source data for the panel b are provided as a Dataset S1.

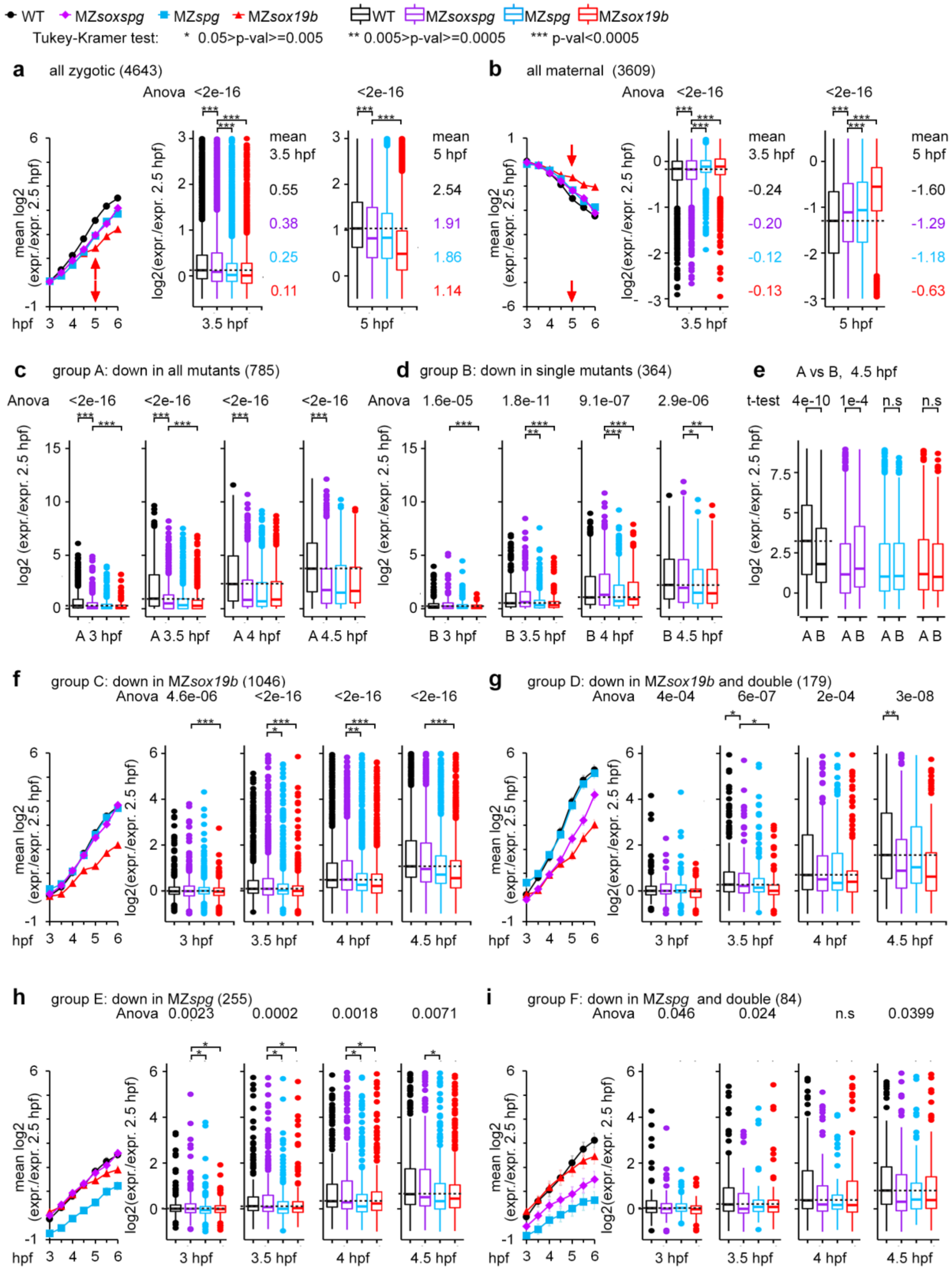

**Figure S5: Non-additive and compensatory effects of Pou5f3 and Sox19b on transcription (supporting information for the main Fig. 3).** Line graphs and box plots are shown for the indicated transcript groups. Line graphs show the mean transcription profiles relative to 2.5 hpf expression at 3 to 6 hpf, in four genotypes (WT, MZ*sox19bspg*, MZ*spg*, MZ*sox19b*). Box plots show the distribution of the relative expression values at the indicated time points for four genotypes (WT, MZ*sox19bspg*, MZ*spg*, MZ*sox19b*). The lower border, middle line and upper border of the box plots correspond to 0.25, 0.5 (median) and 0.75 quantile, respectively Horizontal dashed lines show the WT median level. Statistical significance of the differences between the genotypes were estimated using 1-way Anova. Anova F-values are shown above the graphs, n.s. – non-significant, F >0.05. Pairwise differences for multiple comparisons were estimated by Tukey-Kramer test for (95% confidence level). Significant pairwise differences (p>0.05) between MZ*sox19bspg* and WT, MZ*sox19bspg* and MZ*spg,* and MZ*sox19bspg* and MZ*sox19b* are shown above the graphs (* p<0.05, **p<0.005, ***p<0.0005 in Tukey-Kramer test) in a-d and f-i. All p-values and mean values are listed in the Table S1. **a.** All WT zygotic transcripts. At 3.5 hpf, the zygotic expression in the double mutant was higher than in the single mutants. At 5 hpf, the zygotic expression in the double mutant was the same as in MZ*spg* and higher than in MZ*sox19b*. General delay in the zygotic expression in MZ*sox19b* occurred between 4.5 and 5 hpf, as shown by red arrow. **b.** All WT maternal transcripts. Maternal mRNA levels in the double mutant were lower than in the single mutants at 3.5 hpf and at 5 hpf. General delay in the maternal RNA degradation in MZ*sox19b* occurred between 4.5 and 5 hpf, as shown by red arrow. **c,d.** Non-additive (c, group A) and compensatory (d, group B) effects of Pou5f3 and Sox19b mutations on the earliest zygotic transcription (related to Fig. 3b, d, main text). Box plots show the distribution of zygotic expression levels at 3-4.5 hpf relative to expression at 2.5 hpf. Only significant pairwise differences are shown. WT median - dotted line, p-values in Tukey-Kramer test. **e.** Disbalance of zygotic transcription in MZ*sox19bspg*: A-group transcripts are expressed in a higher level than B-group transcripts in the wild-type, reverse is true for the MZ*sox19bspg*. Box plots for 4.5 hpf. Median - dotted line. p-values in two-tailed Student's t-test, (n.s - p>0.05). **f-i.** Zygotic groups downregulated in the mutants, as indicated (related to the main Fig. 3b, Dataset S1). The differences within the smallest group G (downregulated in the double mutants only) were not significant at any time point (data not shown). MZ*soxspg*=MZ*sox19bspg*. n(wt)=4, n(MZ*spg*)=2, n(MZ*sox19b*)=2, n(double)=2, where n is a number biologically independent experiments. Error bars – standard error of the mean. Source data are provided as a Source Data file and Dataset S1.

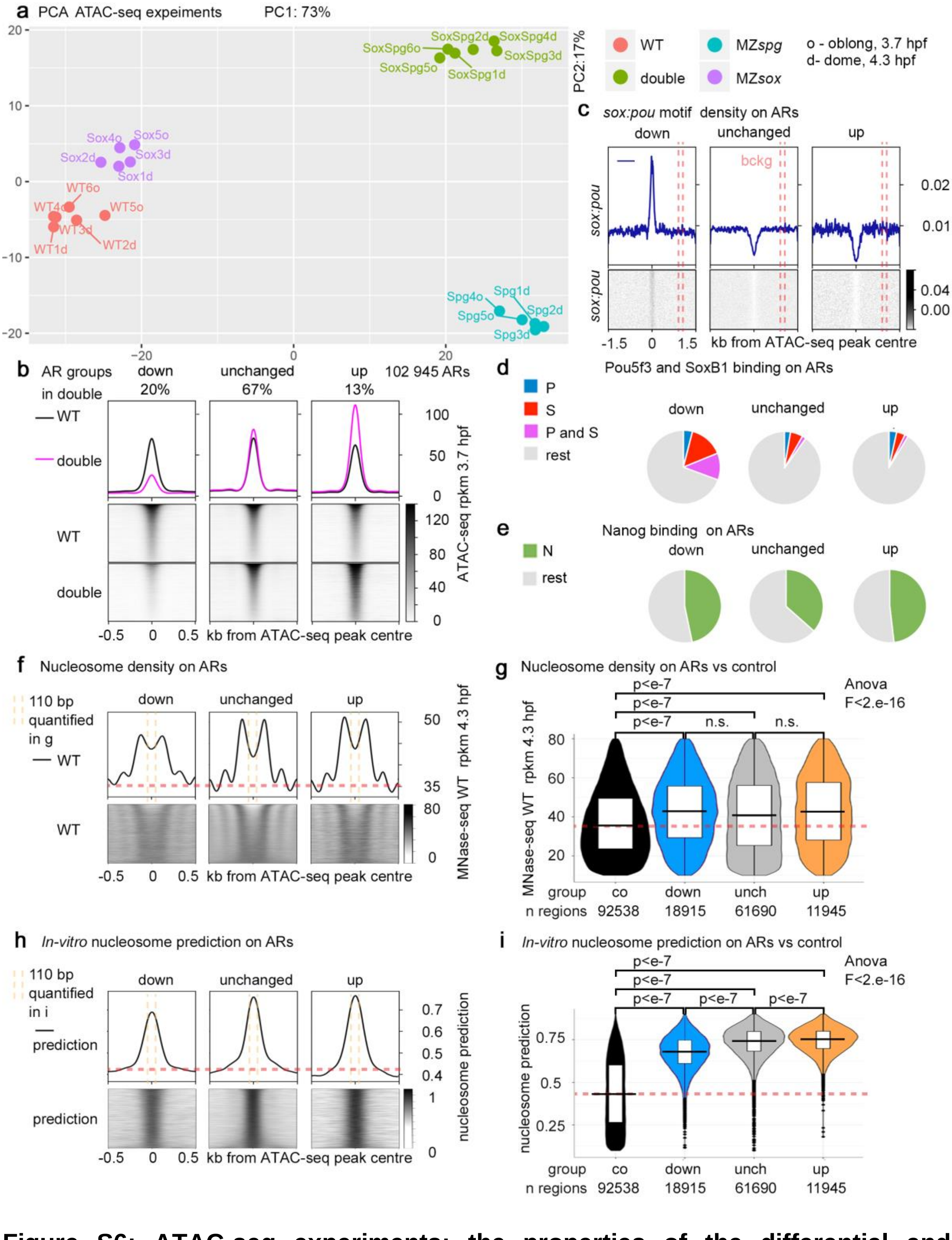

**Figure S6: ATAC-seq experiments; the properties of the differential and unchanged ARs. (supporting information for the main Fig. 5). a.** ATAC-seq experiments for all genotypes were performed at least in triplicates at 4.3 hpf and at least in duplicates at 3.7 hpf. 102945 ATAC-seq peaks present in oblong and dome stages were selected for further analysis. PCA for chromatin accessibility on the selected 102945 ARs in all experiments. Note that the difference between the genotypes exceeds the difference between the time points. We pooled the replicates from two stages to calculate the differences in accessibility between the genotypes. **b**. Chromatin accessibility (3.7 hpf) on three groups of ATAC-seq peaks: reduced (“down”), unchanged, or increased (“up”) in MZ*sox19bspg* double mutants relatively to the wild-type. The regions were sorted by descending ATAC-seq peak score. **c**. *Sox:pou* motifs were enriched in “down” ARs and underrepresented in “unchanged” and “up” ARs. The regions were sorted by descending ATAC-seq peak score. To estimate background motif level, the genomic coordinates of all ATAC-seq peak were shifted 1 kb downstream (red dashed lines). **d**. Overlaps between ARs and ChIP-seq peaks for Pou5f3 and SoxB1[5], at 5 hpf. Note that “down” ARs are 4x more frequently bound by Pou5f3 and SoxB1 than unchanged or “up” ARs (38%, 10% and 8%, respectively). We consider that 38% is likely an underestimated number for TF binding to “down” ARs, because of high cognate motif frequency on these regions (shown in the main Fig. 5f). Also, ChIP-seq was done in a later stage and this technique does not capture all transient TF-binding events. **e.** Overlaps between ARs and ChIP-seq peaks for Nanog at 4.3 hpf stage[6]. Note that “down” and “up” ARs are equally bound by Nanog. Nanog increases chromatin accessibility by displacing nucleosomes[7-9]; presumably, it does so independently on Pou5f3 and Sox19b. **f-i.** Real and predicted nucleosome occupancy on the ATAC-seq peaks (ARs) is higher than genomic mean values (red dashed line). To obtain genomic control values the coordinates of ATAC-seq peaks were shifted 1 kb downstream. **f**. MNase-seq signals at 4.3 hpf (wild-type, n=1 Mnase-seq experiment, data from[9]) were plotted on “down”, “unchanged” and “up” ARs. Note two well-positioned nucleosomes and a gap between them. The 110bp region scored in (g) is in the gap, marked by yellow dashed lines. **g**. MNase-seq signals were quantified on 110 bp around the control, “down” “unchanged” and “up” ARs. Nucleosome occupancy was significantly higher in all ARs than in the control. 1-way Anova, Tukey-Kramer test. The lower border, middle line and upper border of the white box correspond to 0.25, 0.5 (median) and 0.75 quantile, respectively. **h**. *In-vitro* predicted nucleosome occupancy[10] was plotted on “down”, “unchanged” and “up” ARs. The 110-bp region scored in (i) is marked by yellow dashed lines. **i**. *In-vitro* nucleosome prediction values were quantified on 110 bp around the control, “down” “unchanged” and “up” ARs. *In vitro* nucleosome prediction was significantly higher in all ARs than in controls, and significantly differs as “down”<”unchanged”<”up” ARs. The lower border, middle line and upper border of the white box correspond to 0.25, 0.5 (median) and 0.75 quantile, respectively. 1-way Anova, Tukey-Kramer test. Source data are provided as a Dataset S3.

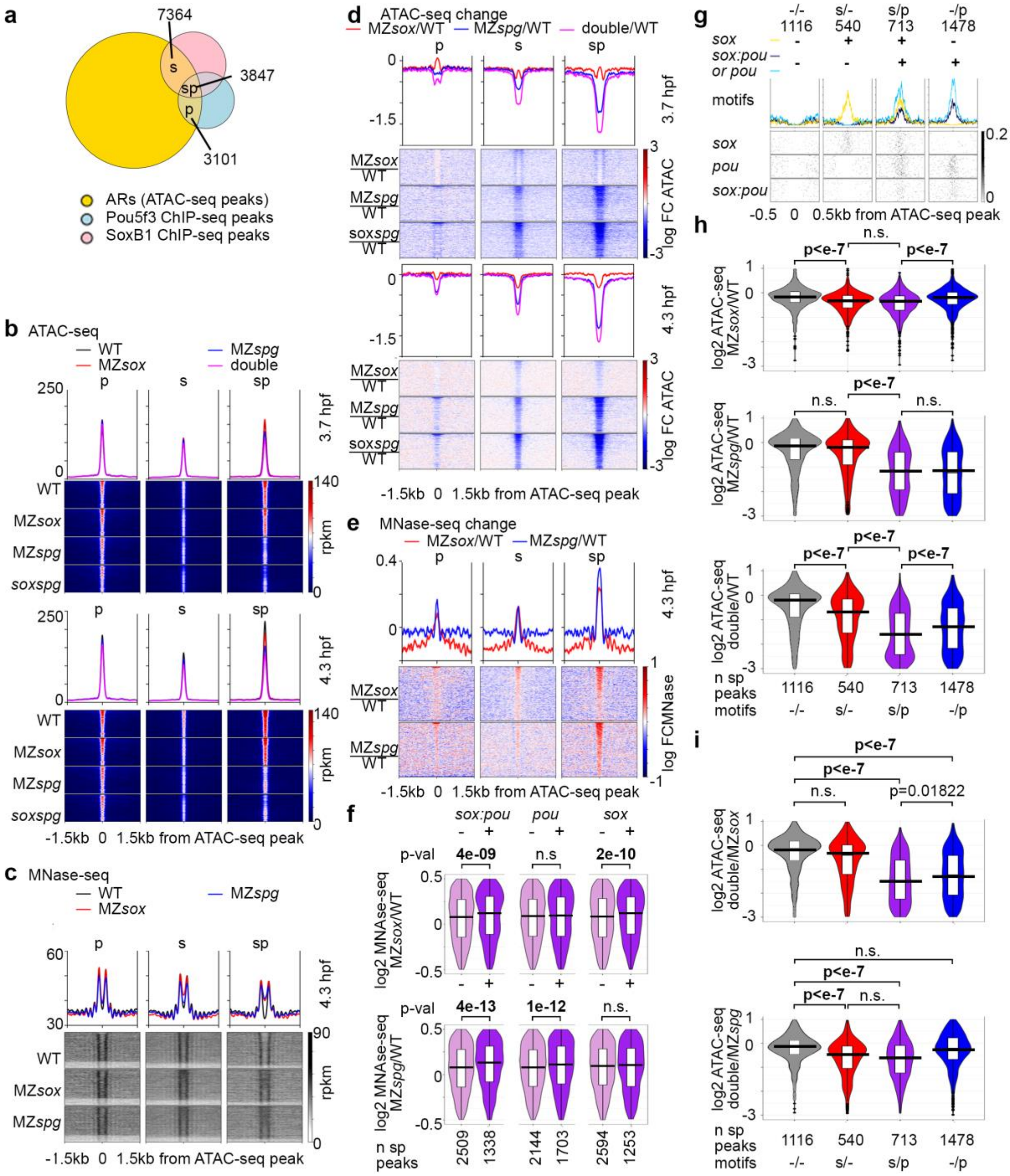

**Figure S7: Sox19b and Pou5f3 promote chromatin accessibility by displacing nucleosomes on different binding motifs (supporting information for the main Fig. 6). a.** Overlap between all ARs (Dataset S3), SoxB1 and Pou5f3 binding peaks[5], (Dataset S4). **b**,**c** ATAC-seq (b) and MNase-seq (c) signals on ARs overlapping Pou5f3-only (p), SoxB1- only (s), or both Pou5f3 and SoxB1 (sp) ChIP-seq peaks in the indicated genotypes. ARs were sorted by descending ATAC-seq peak score. **d.** Differences in chromatin accessibility between the mutants and the wild-type on p, s and sp peaks. Note the strongest effects on sp peaks and in the double mutant MZ*sox19bspg*. **e.** Nucleosome displacement by Pou5f3 and Sox19b on p, s, and sp peaks in the single mutants. Note the strongest effect on sp peaks. **f.** Nucleosome displacement on sp peaks with (+) or without (-) the motifs indicated above was compared. MZ*sox19b* (top): Sox19b displaces nucleosomes on *sox:pou* and *sox*

motifs. MZ*spg* (bottom): Pou5f3 displaces nucleosomes on *sox:pou* and *pou* motifs. Log2 mutant/wt fold changes in MNase signals were calculated per 110-bp regions around AR summits and provided in Dataset S3. P-values in 2-tailed Student t-test. n=1 MNase-seq experiments, WT and MZ*spg* from[9], MZ*sox19b* from this work. **g-i**. Pou5f3 and Sox19b promote chromatin accessibility independently of each other by binding to *pou*/*sox:pou* and *sox* motifs, respectively. n(wt)=3, n(MZ*spg*)=3, n(MZ*sox19b*)=3, n(double)=4, where n is a number biologically independent ATAC-seq experiments. **g**. Chromatin accessibility reduction in the mutants was compared between sp peaks without motifs (-/-), sp peaks with *sox* motifs only (s/-), sp peaks with *sox* and *pou* or *sox:pou* motifs (s/p), or sp peaks with *pou* or *sox:pou* motifs only (-/p). -/- peaks served as control for non-specific effects. Scale- motif density. **h**. Mutants to wild-type change. MZ*sox19b*: chromatin accessibility was reduced to the same extent in s/p and s/- groups, compared to -/- control, indicating that Sox19b promotes accessibility on *sox* motifs independently of Pou5f3 binding nearby. MZ*spg*: chromatin accessibility was reduced to the same extent in s/p and -/p groups, compared to -/- control, indicating that Pou5f3 promotes accessibility on *pou and sox:pou* motifs independently of Sox19b binding nearby. MZ*sox19bspg*: chromatin accessibility was stronger reduced on s/p group compared to s/- and -/p, indicating additive or redundant effects of Pou5f3 and Sox19b independent binding on chromatin accessibility. **i**. Double to single mutant change. MZ*sox19bspg* to MZ*sox19b*: chromatin accessibility is reduced only on peaks with *pou* or *sox: pou* motifs (s/p, -/p) compared to -/- control. MZ*sox19bspg* to MZ*spg*: chromatin accessibility is reduced only on the peaks with *sox* motifs (s/p, s/-) compared to -/- control. We concluded that Pou5f3 and Sox19b promote chromatin accessibility independently. Nucleosome displacing activities of Pou5f3 and Sox19b add up on s/p peaks, as their motifs are located within the same AR. 1-way Anova F-values were <e-16 in all cases. Pairwise differences – Tukey-Kramer test. Mean ATAC-seq values in rkpm were taken for 110 bp around the ATAC-seq peak summit and listed in Dataset S3. double=MZ*sox19bspg*, MZ*sox*=MZ*sox19b.* The lower border, middle line and upper border of the white box in (f,h,i) correspond to 0.25, 0.5 (median) and 0.75 quantile, respectively. Source data are provided as Datasets S3 and S4.

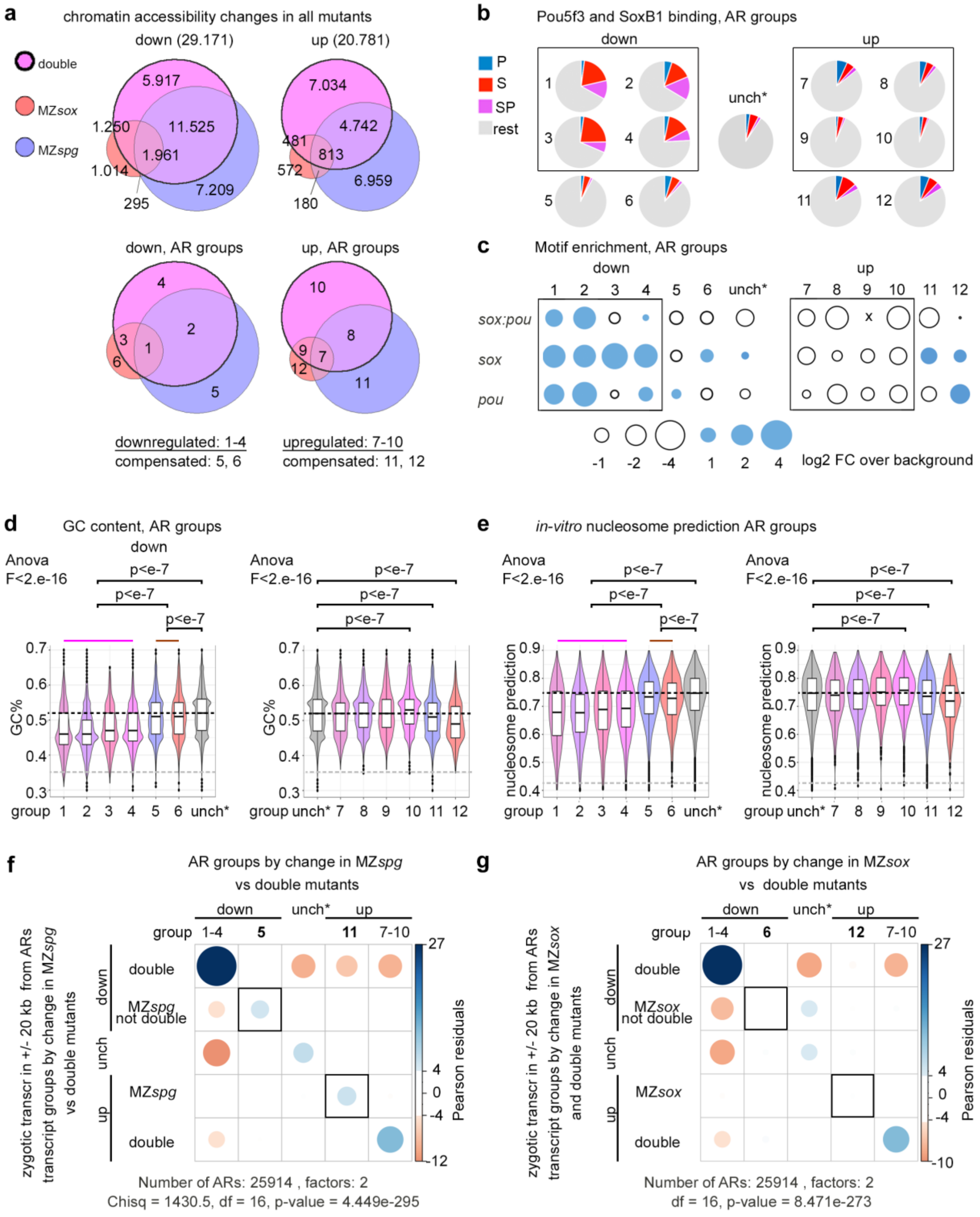

**Figure S8: Properties of ARs with differential chromatin accessibility (supporting information for main Fig. 7). a.** Venn diagrams show overlaps of ARs where chromatin accessibility changed in single and double mutants. The properties of 12 groups were compared in b-e. “Compensated” groups 5, 6, 11, 12 – ARs where chromatin accessibility is changed only in the single mutants and not in double. **b.** Overlap with Pou5f3 and SoxB1 TF binding in 12 groups and unchanged ARs. Note the highest overlap in the groups 1-4. **c**. Motif enrichment in 12 groups and unchanged ARs (x – no motifs). Note, that all the motifs were underrepresented in the upregulated ARs. In compensated ARs, downregulated in single mutant, one type of motifs was underrepresented (except the smallest group 12); this means, that the compensatory effects on chromatin were indirect, *i.e.* not due to the sequence-specific binding of both factors. **d,e.** GC content (d) and *in vitro* predicted nucleosome occupancy in 12 groups and unchanged ARs**.** Note that GC content and *in vitro* predicted nucleosome occupancy is the lowest in the ARs downregulated in the double mutant (gr. 1-4, magenta line) and the highest in the group upregulated in the double mutant only (gr.10). 1-way Anova, Tukey-kramer test. For the “down” groups, the pairwise differences were all significant except 1 and 2, 2 and 4, 3 and 4, 5 and 6. For the “up” groups, groups 7, 8, 9 were not different from each other and from unchanged. Median genomic control level – grey dashed line, median level of unchanged ARs - black dashed line. The lower border, middle line and upper border of the white box correspond to 0.25, 0.5 (median) and 0.75 quantile, respectively. **f,g**. Chi-squared test for independence of transcriptional and chromatin accessibility changes in the single mutants only. P-value is two-tailed. **f.** ARs with down- or upregulated chromatin accessibility in the MZ*spg* are associated with zygotic genes, down- or upregulated in MZ*spg.* **g.** ARs with down- or upregulated chromatin accessibility in the MZ*sox19b* are not significantly associated with zygotic genes, down- or upregulated in MZ*sox19b.* unch* **-** ARs unchanged in both single and double mutants. Source data for the panels a-e are provided as a Dataset S3. Source data for the panel b are provided as a Dataset S4. Source data for the panels f,g are provided as a Dataset S5.

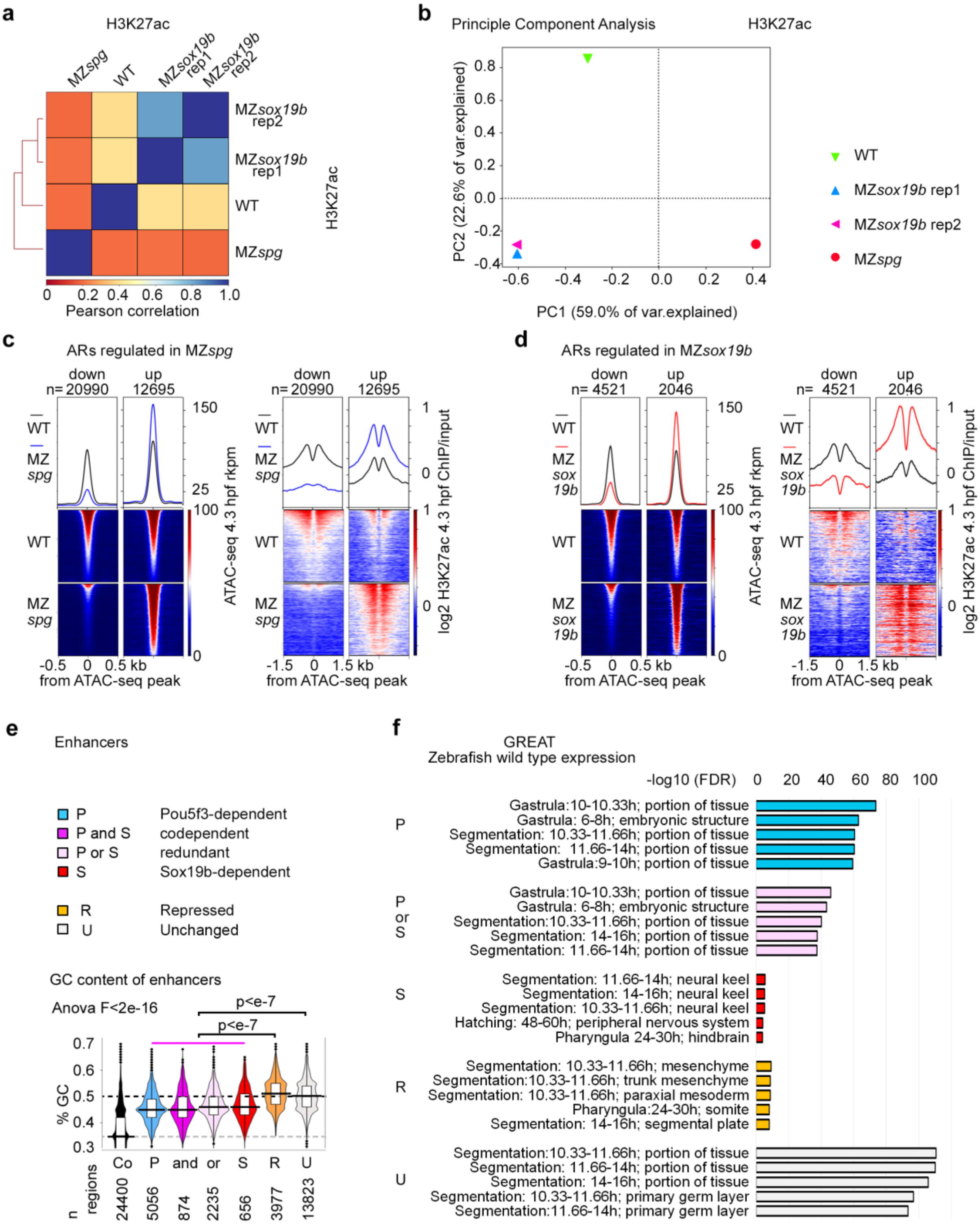

**Figure S9: H3K27ac deposition and chromatin accessibility are coregulated by Pou5f3 and Sox19b (supporting information for main Figures 7 and 8). a-b.** The genome-wide differences between four H3K27ac ChIP-seq experiments: two biological replicates of MZ*sox19b,* and single WT and MZ*spg* replicates were estimated using Pearson correlation (a), and Principle Component Analysis (b). The differences between biological replicates are small compared to the differences between the genotypes. **c-d**. Chromatin accessibility of ARs and H3K27-acetylation of flanking nucleosomes change in parallel in MZ*spg* (c) and MZ*sox19b* (d). ATAC-seq signal (left) and H3K27ac (right) on the downregulated and upregulated ARs in MZ*spg* (c) and

MZ*sox19b* (d). ARs were sorted by descending ATAC-seq peak score. Numbers of regions is shown above the graphs. **e**. Enhancers directly activated by Pou5f3 or Sox19b (Pou5f3-dependent, codependent, redundant and Sox19b-dependent, magenta line) have lower CG content compared to the other enhancers. Gray dashed line – control GC level, black dashed line – GC level of unchanged enhancers. 1-way Anova, Tukey-Kramer test. The lower border, middle line and upper border of the white box correspond to 0.25, 0.5 (median) and 0.75 quantile, respectively. **f.** GREAT[11] enrichment in Gene Ontology term “wild type zebrafish expression” on five indicated enhancer groups. The codependent group (P and S) showed no enrichment. Source data are provided as a Dataset S3.

### Supplementary Table 1: Statistics for the Supplementary Figure 5

| Fig. panel | | Fig. S5a | Fig. S5a | Fig. S5b | Fig. S5b | Fig. S5c | Fig. S5c | Fig. S5c | Fig. S5c |
|---|---|---|---|---|---|---|---|---|---|
| transcript group | | all zygotic 3.5 hpf n=4643 | all zygotic 5 hpf n=4643 | all materna l 3.5 hpf n=3609 | all materna l 5 hpf n=3609 | A 3 hpf n=785 | A 3.5 hpf n=785 | A 4 hpf n=785 | A 4.5 hpf n=785 |
| 1-way Ano va, Pr(>F) | | <2e-16 | <2e-16 | <2e-16 | <2e-16 | <2e-16 | <2e-16 | <2e-16 | <2e-16 |
| Mean log2 (expr/ expr 2.5 h pf) | WT | 0.551 | 2.54 | -0.243 | -1.60 | 0.381 | 1.40 | 2.45 | 3.67 |
| | double | 0.383 | 1.91 | -0.198 | -1.29 | 0.0801 | 0.533 | 1.17 | 1.74 |
| | MZ*spg* | 0.250 | 1.86 | -0.123 | -1.18 | 0.0645 | 0.471 | 1.15 | 1.86 |
| | MZ*sox19b* | 0.115 | 1.14 | -0.129 | -0.631 | -0.182 | 0.196 | 1.03 | 1.84 |
| Tukey multiple compar. of means, 95% conf. level ([1]) | WT-double | 0.0000000 | 0.0000000 | 0.0000010 | 0.0000000 | 0.0000000 | 0.0000000 | 0.0000000 | 0.0000000 |
| | WT-MZ*spg* | 0.0000000 | 0.0000000 | 0.0000000 | 0.0000000 | 0.0000000 | 0.0000000 | 0.0000000 | 0.0000000 |
| | WT-MZ*sox19b* | 0.0000000 | 0.0000000 | 0.0000000 | 0.0000000 | 0.0000000 | 0.0000000 | 0.0000000 | 0.0000000 |
| | double-MZ *spg* | 0.0000000 | *0.6818824* | 0.0000000 | 0.0000530 | *0.9783447* | *0.8581548* | *0.9968089* | *0.8106821* |
| | double-MZ *sox19b* | 0.0000000 | 0.0000000 | 0.0000000 | 0.0000000 | 0.0000000 | 0.0000979 | *0.5753939* | *0.8958933* |
| | MZ*spg*-MZ *sox19b* | 0.0000000 | 0.0000000 | *0.8654144* | 0.0000000 | 0.0000000 | 0.0024627 | *0.7064488* | *0.9977278* |

| panel | | Fig. S5d | Fig. S5d | Fig. S5d | Fig. S5d | Fig. S5f | Fig. S5f | Fig. S5f | Fig. S5f |
|---|---|---|---|---|---|---|---|---|---|
| transcript group | | B 3 hpf n=364 | B 3.5 hpf n=364 | B 4 hpf n=364 | B 4.5 hpf n=364 | C 3 hpf n=1046 | C 3.5 hpf n=1046 | C 4 hpf n=1046 | C 4.5 hpf n=1046 |
| 1-way Ano va, Pr(>F) | | 1.56e-05 | 1.77e-11 | 9.07e-07 | 2.92e-06 | 4.6e-06 | <2e-16 | <2e-16 | <2e-16 |
| Mean log2 (expr/ expr 2.5 h pf) | WT | 0.0760 | 0.682 | 1.48 | 2.51 | 0.0276 | 0.348 | 0.926 | 1.71 |
| | double | 0.0796 | 0.657 | 1.58 | 2.38 | 0.0203 | 0.355 | 0.984 | 1.60 |
| | MZ*spg* | 0.0205 | 0.322 | 0.970 | 1.81 | 0.0312 | 0.243 | 0.757 | 1.42 |
| | MZ*sox19b* | -0.106 | 0.117 | 0.920 | 1.70 | -0.0525 | 0.0416 | 0.438 | 0.969 |
| Tukey multiple compar. of means, 95% conf. level ([1]) | WT-double | *0.9997748* | *0.9923292* | *0.9188907* | *0.9022907* | *0.9795195* | *0.9979221* | *0.8057498* | *0.5238130* |
| | WT-MZ*spg* | *0.5536120* | 0.0004525 | 0.0034788 | 0.0008908 | *0.9973472* | 0.0338885 | 0.0411882 | 0.0014865 |
| | WT-MZ*sox19b* | 0.0001062 | 0.0000000 | 0.0010069 | 0.0000798 | 0.0000912 | 0.0000000 | 0.0000000 | 0.0000000 |
| | double-MZ *spg* | *0.4992449* | 0.0013706 | 0.0002894 | 0.0103997 | *0.9357449* | 0.0200573 | 0.0022833 | *0.0997061* |
| | double-MZ *sox19b* | 0.0000724 | 0.0000000 | 0.0000679 | 0.0013812 | 0.0004911 | 0.0000000 | 0.0000000 | 0.0000000 |
| | MZ*spg*-MZ *sox19b* | 0.0151644 | *0.1112940* | *0.9873325* | *0.9400947* | 0.0000372 | 0.0000013 | 0.0000045 | 0.0000002 |

| panel | | Fig. S5g | Fig. S5g | Fig. S5g | Fig. S5g | Fig. S5h | Fig. S5h | Fig. S5h | Fig. S5h |
|---|---|---|---|---|---|---|---|---|---|
| transcript group | | D 3 hpf n=179 | D 3.5 hpf n=179 | D 4 hpf n=179 | D 4.5 hpf n=179 | E 3 hpf n=255 | E 3.5 hpf n=255 | E 4 hpf n=255 | E 4.5 hpf n=255 |
| 1-way Ano va, Pr(>F) | | 0.000456 | 6.06e-07 | 0.00026 | 3.49e-08 | 0.00228 | 0.000201 | 0.00185 | 0.00714 |
| Mean log2 (expr/ expr 2.5 h pf) | WT | 0.0917 | 0.750 | 1.50 | 2.54 | 0.105 | 0.512 | 0.983 | 1.43 |
| | double | 0.0226 | 0.422 | 1.03 | 1.58 | 0.133 | 0.517 | 1.05 | 1.42 |
| | MZ*spg* | 0.115 | 0.482 | 1.27 | 2.12 | -0.00753 | 0.215 | 0.590 | 0.933 |
| | MZ*sox19b* | -0.116 | 0.0831 | 0.686 | 1.18 | 0.00826 | 0.232 | 0.662 | 1.09 |
| Tukey multiple compar. of means, 95% conf. level ([1]) | WT-double | *0.6516316* | 0.0283082 | *0.0749138* | 0.0003358 | *0.9291132* | *0.9999295* | *0.9708215* | *0.9997467* |
| | WT-MZ*spg* | *0.9807283* | *0.1051676* | *0.6477133* | *0.2875473* | *0.0561773* | 0.0075374 | 0.0322962 | 0.0216611 |
| | WT-MZ*sox19b* | 0.0029939 | 0.0000001 | 0.0001794 | 0.0000001 | *0.1308687* | 0.0135442 | *0.1160139* | *0.1970007* |
| | double-MZ *spg* | *0.4110574* | *0.9570867* | *0.5957786* | *0.1051110* | 0.0094600 | 0.0062037 | 0.0083271 | 0.0280911 |
| | double-MZ *sox19b* | *0.0949162* | 0.0218842 | *0.2841881* | *0.3083827* | 0.0279291 | 0.0112710 | 0.0382023 | *0.2334347* |
| | MZ*spg*-MZ *sox19b* | 0.0007044 | 0.0042592 | 0.0135595 | 0.0003983 | *0.9848318* | *0.9978702* | *0.9581222* | *0.8081560* |

1

[1] Non-significant p-values (P>0.05) are in italics

**Supplementary Table S2. Data Resources (published and generated during this work)**

| REAGENT or RESOURCE | SOURCE | IDENTIFIER |
|---|---|---|
| Antibodies | | |
| Anti-Histone H3 (acetyl K27) rabbit, 1/100 dilution | Abcam plc., Cambridge, UK | ab 4729 |
| Anti-Histone H3 (tri-methyl K4) rabbit, 1/100 dilution | Millipore Co., Temecula, California, USA | 07-449 |
| Bacterial and Virus Strains | | |
| One Shot™ TOP10 chemically competent *E. coli* | Invitrogen™ | C404003 |
| Chemicals, Peptides, and Recombinant Proteins | | |
| cOmplete™, EDTA-free Protease Inhibitor Cocktail | Sigma-Aldrich Chemie GmbH, Germany | 5056489001 |
| Micrococcal Nuclease | Sigma-Aldrich Chemie GmbH, Germany | N3755-200UN |
| SYTOX Green | ThermoFisher SCIENTIFIC | S7020 |
| Critical Commercial Assays | | |
| Agencourt® AMPure® XP Beads | Beckmann Coulter, Krefeld, Germany | A63880 |
| Agilent High Sensitivity DNA Kit | Agilent Technologies, Santa Clara, California, USA | 5067-4626 |
| Agilent RNA 6000 Nano Kit | Agilent Technologies, Santa Clara, California, USA | 5067-1511 |
| Dynabeads® Protein G | invitrogen Dynal AS, Oslo, Norway | 10003D |
| E.Z.N.A® Cycle Pure Kit | Omega Biotek, Norcross, Georgia, USA | D6493-02 |
| Microcon®-30 Centrifugal Filters | Merck Millipore, Darmstadt, Germany | MRCF0R030 |
| MinElute® PCR Purification Kit (50) | Qiagen GmbH, Hilden, Germany | 28004 |
| mMESSAGE mMACHINE® SP6 transcription Kit | Ambion | 10086184 |
| NEBNext® High Fidelity Master Mix | New England biolabs Inc. | M0541L |
| NEBNext Ultra DNA Library Prep Kit for Illumina | New England Biolabs, Inc., Frankfurt a.M., Germany | E7370S |
| NEBNext® Multiplex Oligos for Illumina® (Index Primers Set 1) | New England Biolabs, Inc., Frankfurt a.M., Germany | E7335S |
| RNeasy® Mini Kit | QIAGEN, Hilden, Germany | 74104 |
| Quant-iT™ PicoGreen® dsDNA Assay Kit | invitrogen™ Molecular Probes® Waltham, Massachusetts, USA | Q33120, P11496 |
| SPRIselect | Beckman Coulter Inc., Brea, California, USA | B23317 |
| Tagment DNA Enzyme and Buffer (Small kit) | Illumina, Inc. San Diego, California, USA | 20034210 |

| | | |
|---|---|---|
| Deposited Data | | |
| ATAC-seq of the WT, MZ*sox19b*, MZ*spg* and MZ*sox19bspg*, 3.7 hpf and 4.3 hpf zebrafish embryos | this work | GEO: GSE188364 |
| ChIP-seq for H3K27ac and H3K4me3 in three genotypes, 4.3 hpf | this work | GEO: GSE143306 |
| ChIP-seq for Pou5f3 and SoxB1, 5 hpf | Leichsenring et al., 2013 | GEO: GSE39780 |
| Consensus motifs enriched in accessible regions | This work | Supplementary Table S4 |
| List of Pou5f3, SoxB1 and Nanog ChIP-seq peaks | Data from Leichsenring et al., 2013, and Xu et al., 2011, remapped to GRCz11 in this work | Supplementary Table S4 |
| MNase-seq of MZ*sox19b* embryos, 4.3 hpf | this work | GEO: GSE125945 |
| MNase-seq of WT and MZ*spg* embryos, 4.3 hpf | Veil et al., 2019 | GEO: GSE109410 |
| Time-resolved RNA-seq at 8 time points (2.5 hpf to 6 hpf, 30 min intervals) in the wild-type, MZ*spg*, MZ*sox19b* and MZ*sox19bspg* embryos | this work | GEO: GSE137424, Supplementary Tables S1 and S2 |
| ChIP-seq for Pou5f3 and SoxB1 | Leichsenring et al., 2013 | GEO: GSE39780 |
| ChIP-seq for Nanog | Xu et al., 2011 | GEO: GSE34683 |
| Zebrafish reference genome assembly danrer11/ GRCz11 | Genome reference Consortium | https://www.ncbi.nlm.nih.gov/grc/zebrafish |
| Experimental Models: Organisms/Strains | | |
| Wild-type zebrafish strain AB/TL | ZIRC | ZL1/ZL86 |
| MZ*spg* zebrafish | Lunde et al., 2004 | m793 |
| MZ*sox19b* zebrafish | this work | m1434 |
| Oligonucleotides | | |
| MO3-Sox2 Sox2 Morpholino 5' - GAAAGTCTACCCCACCAGCCGTAAA - 3' | Okuda et al., 2010 | ZFIN ID: ZDB-MRPHLNO-080329-1 |
| MO4-Sox2 Sox2 Morpholino 5' - GAGAGGCTGCTGAAGTTACCTTAGC - 3' | Okuda et al., 2010 | ZFIN ID: ZDB-MRPHLNO-080329-2 |
| MO3-Sox3 Sox3 Morpholino 5' - TACATTCTTAAAAGTGGTGCCAAGC - 3' | Okuda et al., 2010 | ZFIN ID: ZDB-MRPHLNO-100527-3 |
| MO4-Sox3 Sox3 Morpholino 5' - GAAGTCAGTCAAAAGTTCAGAGAGC - 3' | Okuda et al., 2010 | ZFIN ID: ZDB-MRPHLNO-100527-4 |
| MO1-Sox19a Sox19a Morpholino 5' - GTACATGGCTGCCAACAGAAGTTAG - 3' | Okuda et al., 2010 | ZFIN ID: ZDB-MRPHLNO-100527-5 |
| MO2-Sox19a Sox19a Morpholino 5' - AAAACGAGAGCGAGCCGTCTGTAAC - 3' | Okuda et al., 2010 | ZFIN ID: ZDB-MRPHLNO-100527-6 |
| MO1-Sox19b Sox19b Morpholino 5' - GTACATCATGCCACTTCTCGCTTTG - 3' | Okuda et al., 2010 | ZFIN ID: ZDB-MRPHLNO-100527-7 |
| MO2-Sox19b Sox19b Morpholino 5' - ACGAGCGAGCCTAATCAGGTCAAAC - 3' | Okuda et al., 2010 | ZFIN ID: ZDB-MRPHLNO-100527-8 |
| MOa-nog1 Noggin1 Morpholino 5' - GCGGGAAATCCATCCTTTTGAAATC - 3' | Dal-Pra et al., 2006 | ZFIN ID: ZDB-MRPHLNO-080212-1 |

| | | |
|---|---|---|
| MO1-chrd Chordin Morpholino 5' -ATCCACAGCAGCCCCTCCATCATCC - 3' | Dal-Pra et al., 2006 | ZFIN ID: ZDB-MRPHLNO-050221-6 |
| Index 1 (i7) Primer N701 CAAGCAGAAGACGGCATACGAGATTCGCCTTAGTCTCGTGGGCTCGG | Illumina Nextera DNA Index | |
| Index 1 (i7) Primer N702 CAAGCAGAAGACGGCATACGAGATCTAGTACGGTCTCGTGGGCTCGG | Illumina Nextera DNA Index | |
| Index 1 (i7) Primer N703 CAAGCAGAAGACGGCATACGAGATTTCTGCCTGTCTCGTGGGCTCGG | Illumina Nextera DNA Index | |
| Index 1 (i7) Primer N704 CAAGCAGAAGACGGCATACGAGATGCTCAGGAGTCTCGTGGGCTCGG | Illumina Nextera DNA Index | |
| Index 1 (i5) Primer N501 AATGATACGGCGACCACCGAGATCTACACTAGATCGCTCGTCGGCAGCGTC | Illumina Nextera DNA Index | |
| Index 1 (i5) Primer N502 AATGATACGGCGACCACCGAGATCTACACCTCTCTATTCGTCGGCAGCGTC | Illumina Nextera DNA Index | |
| Index 1 (i5) Primer N503 AATGATACGGCGACCACCGAGATCTACACTATCCTCTTCGTCGGCAGCGTC | Illumina Nextera DNA Index | |
| Index 1 (i5) Primer N504 AATGATACGGCGACCACCGAGATCTACACAGAGTAGATCGTCGGCAGCGTC | Illumina Nextera DNA Index | |
| PCR primer for genotyping *sox19b* mutants Sox19b-f1 5'-ATTTGGGGTGCTTTCTTCAGC-3' | this work | no |
| PCR primer for genotyping *sox19b* mutants Sox19b-r1 5'-GTTCTCCTGGGCCATCTTCC-3' | this work | no |
| sox19bF1: Sox19b PCR cloning primer<br><br>sox19Bf1 with BamHI: 5'-GGGGGATCCATGGAGCACGAGCTGAAGAC-3', | this work | no |
| Sox19b PCR cloning primer<br><br>Sox19bR1 with Xho1: 5'-GGGGCTCGAGTCAGATGTGAGTGAGGGGAAC-3' | this work | no |
| PCR primer for genotyping *spg* mutants spg-f1 GTCGTCTGACTGAACATTTTGC | this work | no |
| PCR primer for genotyping *spg* mutants spg-r1 GCAGTGATTCTGAGGAAGAGGT | this work | no |
| PCR primer for ChIP-seq control, positive reference tiparp_f_1 5' CGCTCCCAACTCCATGTATC-3' | this work | no |
| PCR primer for ChIP-seq control, positive reference tiparp_r_1 5'-AACGCAAGCCAAACGATCTC-3' | this work | no |
| PCR primer for ChIP-seq control, negative reference igsf2_f_2 5'-GAACTGCATTAGAGACCCAC-3' | this work | no |
| PCR primer for ChIP-seq control, negative reference igsf2_r_2 5'-CAATCAACTGGGAAAGCATGA-3' | this work | no |
| Recombinant DNA | | |
| CS2+Sox19b plasmid for mRNA synthesis | this work | no |
| Software and Algorithms | | |

| | | |
|---|---|---|
| Bed Tools | Quinlan and Hall,2010 | BED Tools in usegalaxy.eu |
| Bowtie2 | Langmead and Salzberg, 2012 | Bowtie2 in usegalaxy.eu |
| DeepTools2 | Ramirez et al., 2016 | deepTools in usegalaxy.eu |
| DESeq2 | Love et al., 2014 | DESeq2 in usegalaxy.eu |
| FeatureCounts | Liao et al., 2014 | featureCounts in usegalaxy.eu |
| Galaxy server | Afgan et al., 2018 | https://usegalaxy.eu/ |
| GREAT: Genomic Regions Enrichment of Annotations Tool, version 3.0.0 | Hiller et al., 2013 | http://great.stanford.edu/great/public-3.0.0/html/ |
| *In-vitro* nucleosome prediction program | Kaplan et al., 2009 and https://github.com/bgruening/galaxytools | Nucleosome Predictions in usegalaxy.eu |
| MACS2 | Ferg et al., 2007 | MACS2 callpeak and MACS2 bdgpeakcall in usegalaxy.eu |
| R programming packages | see the complete list in the Supplementary data | see the complete list in the Supplementary data |
| RNA Star | Dobin et al., 2013 | RNA Star in usegalaxy.eu |
| RNA-sense | this work | https://bioconductor.org/packages/release/bioc/html/RNAsense.html |

#### Supplementary References.

1 Okuda, Y., Ogura, E., Kondoh, H. & Kamachi, Y. B1 SOX coordinate cell specification with patterning and morphogenesis in the early zebrafish embryo. *PLoS Genetics* **6**, e1000936, doi:10.1371/journal.pgen.1000936 (2010).

2 Harvey, S. A. *et al.* Identification of the zebrafish maternal and paternal transcriptomes. *Development* **140**, 2703-2710, doi:10.1242/dev.095091 (2013).

3 Haberle, V. *et al.* Two independent transcription initiation codes overlap on vertebrate core promoters. *Nature* **507**, 381-385, doi:10.1038/nature12974 (2014).

4 Lee, M. T. *et al.* Nanog, Pou5f1 and SoxB1 activate zygotic gene expression during the maternal-to-zygotic transition. *Nature*, 360-364., doi:10.1038/nature12632 (2013).

5 Leichsenring, M., Maes, J., Mossner, R., Driever, W. & Onichtchouk, D. Pou5f1 transcription factor controls zygotic gene activation in vertebrates. *Science* **341**, 1005-1009, doi:10.1126/science.1242527 (2013).

6 Xu, C. *et al.* Nanog-like Regulates Endoderm Formation through the Mxtx2-Nodal Pathway. *Developmental cell* **22**, 625-638, doi:10.1016/j.devcel.2012.01.003 (2012).

7 Miao, L. *et al.* Synergistic activity of Nanog, Pou5f3, and Sox19b establishes chromatin accessibility and developmental competence in a context-dependent manner. *bioRxiv*, 2020.2009.2001.278796, doi:10.1101/2020.09.01.278796 (2020).

8 Palfy, M., Schulze, G., Valen, E. & Vastenhouw, N. L. Chromatin accessibility established by Pou5f3, Sox19b and Nanog primes genes for activity during zebrafish genome activation. *PLoS Genet* **16**, e1008546, doi:10.1371/journal.pgen.1008546 (2020).

9 Veil, M., Yampolsky, L., Gruening, B. & Onichtchouk, D. Pou5f3, SoxB1, and Nanog remodel chromatin on High Nucleosome Affinity Regions at Zygotic Genome Activation. *Genome Res*, doi:10.1101/gr.240572.118 (2019).

10 Kaplan, N. *et al.* The DNA-encoded nucleosome organization of a eukaryotic genome. *Nature* **458**, 362-366, doi:10.1038/nature07667 (2009).

11 Hiller, M. *et al.* Computational methods to detect conserved non-genic elements in phylogenetically isolated genomes: application to zebrafish. *Nucleic Acids Res* **41**, e151, doi:10.1093/nar/gkt557 (2013).
